# Kin discrimination in plants: competitive ability over kinship response

**DOI:** 10.64898/2026.07.24.740458

**Authors:** Lucas Mazal, Philippe Malagoli, Amélia Bourceret, Marc-André Selosse, Dov Corenblit, Irène Till-Bottraud, Boris Fumanal

**Affiliations:** Université Clermont Auvergne, CNRS, GEOLAB, F-63000 Clermont-Ferrand, France; VetAgro Sup, INRAE UMR 0874 UREP, Université Clermont Auvergne, Clermont-Ferrand, France; Université Clermont Auvergne, INRAE, UMR 547 PIAF, F-63000 Clermont-Ferrand, France; Institut de Systématique, Évolution, Biodiversité (ISYEB), Muséum national d’Histoire naturelle, CNRS, Sorbonne Université, EPHE, Université des Antilles, Paris, France; Department of Plant Taxonomy and Nature Conservation, University of Gdansk, Wita Stwosza 59, 80-308, Gdansk, Poland; Institut Universitaire de France; Laboratoire écologie fonctionnelle et environnement, Université Paul Sabatier, 118 Route de Narbonne, Toulouse 31062, CEDEX 9, France

**Keywords:** Kin recognition, Intraspecific interactions, Competitive ability, Resource partitioning, Black poplar

## Abstract

Numerous studies on plant–plant interactions aim to distinguish positive from negative effects, especially among related individuals. Most studies interpret a reduction in biomass when grown with related individuals as kin recognition. However, these interactions can shift from positive to negative depending on environmental or biotic factors. We investigated these interactions in Black poplar (*Populus nigra* L.), a riparian tree species growing in high densities and stressful habitats, by examining the effects of relatedness and water deficit during early developmental stages. We set up two experiments where seedlings from four kin groups were competing with related or unrelated individuals, in well-watered versus water deficit environment, and studied their response through growth, root nitrogen uptake and mycorrhizal diversity. The four kin groups exhibit similar intrinsic growth capacities. However, for two kin groups, we found differences in competitive ability. Thus, we found that individuals responded differently depending on whether they were grown with related or unrelated individuals. Water deficit did not change the outcomes of interactions. Differences in competitive ability between kin groups was the most parsimonious interpretation, thus eliminating kin recognition as an explanation. Our study suggests that results obtained in general in kin studies must be carefully analyzed and that alternative interpretations, such as differences in competitive ability, must be tested before conclude wrongly to kin recognition. Our study also suggests that mycorrhizal communities develop differently according to the kin groups considered and can potentially have impacts on the growth of individuals.

## Introduction

As terrestrial plants are fixed to their substrate, they are in contact with their direct neighbours throughout their lifetime. Notably, some plants seem to discriminate their conspecific neighbours based on their genetic relatedness (kin recognition) and can adapt their ’behaviour’ accordingly (kin discrimination) (Anten and Chen, 2021; Ehlers and Bilde, 2019; Mazal et al. 2023). Kin recognition, defined as the ability to recognize related individuals could be advantageous in reducing competition with them (West et al. 2007; File et al. 2012a), which amounts to a cooperative behaviour that may be promoted by kin selection. The theoretical framework behind kin recognition and discrimination is kin selection theory (Hamilton 1964), which states that altruistic phenotypes are favored when the cost for the actor (-c) is lower than the benefit received by the recipient (b) weighted by the relatedness (r) between the actor and the recipient (-c < br).

In practice, the mechanisms of kin recognition are not easily observable and studies mainly focus on kin discrimination which corresponds to the observation of differences in the expression of traits of individuals when grown with kin or with strangers (Mazal et al. 2023). The confusion between kin recognition and kin discrimination probably comes from the fact that the preferred way to detect kin recognition in plants is through its expression (trait variation between kin and non-kin neighbour treatments) which translates to kin discrimination. The last decade, numerous studies have highlighted kin interactions in plants, especially kin discrimination for exploitation of soil or light resources. In previous reviews, Anten and Chen (2021) and Mazal et al. (2023) provide a critical and detailed overview of kin discrimination in plants. Although there are various methodological and interpretation biases in these studies (see also Karst et al. 2023), the current state of knowledge seems to point to the existence of kin recognition and discrimination in plants (Anten and Chen, 2021; Biedrzycki and Bais, 2022; Buwa and Pannell, 2022). A common finding is that plants growing with related individuals exhibit a reduction in belowground and aboveground biomass compared to plants growing with non-related neighbours (Anten and Chen, 2021), which is interpreted as a reduction in competition towards relatives, possibly leading to positive interactions between related individuals (File et al. 2012a). In another review, Biedrzycki and Bais (2022) investigated studies based on nutrient and resource allocation and discussed how plant roots are involved in kin interactions processes. Similarly to what is observed for biomass, several studies found that plants growing with kin reduce their nutrient uptake, especially nitrogen (Zhang et al. 2016; Li et al. 2018; Pezzola et al. 2019). Because plants can modulate their nutrient niches based on their efficiency in nutrient uptake, it has been suggested that niche differentiation can lead to species coexistence by lessening competition (McKane et al. 2002; Xu et al. 2016). Therefore, studying nutrient niches could be a possible cue to detect kin recognition in plants (Li et al. 2018) and in particular, nitrogen has been proposed as an ideal nutrient to examine root interactions when plants of the same species are grown together (Zhang et al. 2016). However, interactions (including kin interactions) can be affected in predictable ways by current ecological conditions (“conditionality” concept from Bronstein 1994). For terrestrial plants, drought is an important abiotic stress factor that can strongly modify belowground interactions (reviewed by Foxx and Fort, 2019). Under water deficit, root allocation increases, and rooting depth may also increases (Jackson and Colmer, 2005) which will modify root interactions between neighbours. Plant root systems must optimize the uptake of several limiting resources, and it is well established that the intensity of belowground interactions varies with the availability of soil resources (de Vries and Archibald, 2018). Thus, interactions between plants may be modified in stressful or resource-limited environments (Litav and Harper, 1967). The stress gradient hypothesis predicts that different stress levels will change the type of interactions between individuals from positive (i.e. facilitation) in stressful environments to negative (i.e. competition) in favourable environments (Bertness and Callaway, 1994, Callaway et al. 2002). In several species, kin recognition responses were triggered by stress such as nutrient availability (Li et al. 2018; Pezzola et al. 2020) and heavy metal concentrations (Li et al. 2018). However, water availability did not trigger a particular response toward related individuals in *Glechoma hederacea* (Goddard et al. 2020).

In addition, most plants establish symbiotic associations with soil fungi, the so-called mycorrhizas (Smith & Read, 2008; van der Heijden et al., 2015), where they exchange carbon for mineral nutrients collected in soil. These fungi can be shared between neighbouring plants, forming common mycorrhizal networks (CMN; Selosse et al. 2006; Karst et al. 2023), which have been suggested to play a role in kin recognition (Anten and Chen, 2021; Biedrzycki and Bais, 2022). In fact, individuals can exchange nutrients if they are connected by CMN, in particular carbon (Gonneau et al. 2014; Merckx 2013), phosphorus (Whiteside et al. 2019), nitrogen (He et al. 2003), water (Bingham and Simard, 2011), and likely other compounds, such as defence signals (Song et al. 2015), with each other. It has been suggested that relatedness between two individuals is a key factor in the establishment of a mycorrhizal network but also in communication between individuals, and in the exchange of organic compounds between plants (Dudley et al. 2015; Pickles et al. 2017; Tedersoo et al. 2020). Root exudates have also been proposed as mediators of kin recognition in *Deschampsia caespitosa* (Semchenko et al. 2014) and in *Oryza sativa* (Yang et al. 2018). Moreover, CMN involving ectomycorrhizal fungi, the mycorrhizal fungi dominating in temperate forests, have been demonstrated to transfer water, nitrogen, and small quantities of carbon between Douglas-fir hosts (Pickles et al. 2017). Despite some bias in the literature, such as the lack of analysis of confounding effects (Karst et al. 2023), evidence that closely related plants display greater CMN size and root colonization (File et al. 2012b; Dudley et al. 2013) raises the possibility of preferential connectivity between related individuals especially in trees (Pickles et al. 2017; Tedersoo et al. 2020). If that is the case, relatedness may influence nutrient uptake or transfer by altering root growth, hence mycorrhizal and the level of fungal sharing (CMN).

*Populus nigra* L. (Salicaceae) is a dioicous pioneer riparian tree species occurring along many European river courses (Villar and Forestier, 2006) with a good tolerance to submersion, sediment burial and high temperatures (Chamaillard 2011). *P. nigra* is more or less drought avoidant, or even tolerant for some genotypes (Garavillon-Tournayre et al. 2018). Moreover, from the early stages of development, *P. nigra* forms a dense and extensive rhizosphere (Gardes et al. 2003) that interacts with endo- and mostly ectomycorrhizal fungal species (Gryta et al. 2006; Doty et al. 2009; Harner et al. 2011). During their first development stages, young black poplar individuals experience stressful conditions such as drought, submersion and strong mechanical constrains (Vautier et al. 2016), which, combined with high seedling densities, enhancing competition and facilitation (Corenblit et al. 2009; Hortobágyi et al. 2018). Facilitation is ‘an interaction in which the presence of one species alters the environment in a way that enhances the growth, survival and reproduction of a second species’ (Bronstein, 2009), a definition also valid for different genotypes within a species (McIntire and Fajardo, 2014). For *P. nigra*, facilitation is enhanced by niche construction: individuals trap sediments which result in a massive deposit of sediments and organic matter. Mazal et al. (2021) found a significant spatial genetic structure in youngest *P. nigra* cohorts (5 years-old) on alluvial bars, indicating that related individuals grow close to each other. Interactions between related individuals may take place early in the life cycle. Positive intraspecific interactions are suspected to be a major driver at the establishment stage (i.e. first 5 years) in dense recruitment patches of *P. nigra* (Barsoum 2002; Corenblit et al. 2014, 2018).

The present study investigated the intraspecific interactions between young black poplars in early-life stages in relation to relatedness with their neighbours and to a water deficit. We address the following questions: (1) are black poplars capable of kin discrimination, i.e. do they show different responses (growth traits) according to neighbour’s genetic identity (kin *versus* strangers)? (2) does a drought modulate the outcome of these interactions? (3) does relatedness between individuals influence the diversity of the associated CMN? and finally (4) is root nutrient uptake affected by neighbour’s identity? To do so, we set up two different experiments where individual black poplar seedlings were competing with related individuals (kin) or unrelated individuals (strangers) in a non-stressful or stressful environment. We studied growth, root-associated fungal communities, with a focus on nitrogen uptake and mycorrhizal diversity.

## Materials and Methods

### Plant material

Seeds of *P. nigra* were collected in April 2019 from one population of the Allier river: four mother trees along the Allier River, named A1, A2, A3, A4. Mother trees were separated by a minimum of 2 kms to minimize the risk of collecting seeds sired by the same father tree. Seeds from the same mother tree are either half-sibs (same mother but different fathers) or full-sibs (same mother and father) and are considered a family. Seeds were extracted from the seedpods manually, separated from their cotton fluff and stored at -18°C until sowing.

### Experimental setup

We setup two experiments (experiment #1 and #2 in the following sections) in each of which seeds were arranged in 2L pots (12x12x20cm) with the following kinship treatments: Kin treatment, a focal plant surrounded by three neighbours from the same family; Stranger treatment, a focal plant surrounded by plants belonging to each of the three other families. In the Kin and Stranger treatments plants were placed at two centimetres from the focal plant. The sifted sand substrate used in our pots was collected directly from the riverbanks of the Allier River near Mirefleurs village (45°41’24.1” N 3°12’09.7” E). Seeds from each family were germinated in June 2019 in a pot filled with the same sifted sand. One to two-days-old seedlings were selected for uniformity in size and transplanted into the experimental pots. Under each pot, an individual plastic dish prevented seedling dehydration. Pots from the same experiment were randomly placed in a greenhouse compartment at the Clermont Auvergne University (natural light, temperature and humidity maintained at +25°C and 60% respectively). All the pots were randomized in the compartment every two months. The experiments lasted one year, from June 2019 to June 2020.

### Experiment#1 Kin discrimination in water-deficit condition

This experiment aimed to answer the following questions: (1) are black poplars capable of kin discrimination, i.e. do they show different responses (growth traits) according to neighbour’s genetic identity (kin *versus* strangers)? (2) does a water deficit modulate the outcome of these interactions? To answer these questions, we use the kinship treatments (Kin *versus* Stranger, explained above in the *Experimental setup* section) and in addition a Control treatment was set up with one focal plant alone per pot for each family. The water-deficit condition was obtained by watering pots at 20% of the field capacity versus 100% for well-watered treatment. In total, for each of the four families, each treatment (Kin, Stranger, Control) was replicated six times in both the water deficit and well-watered treatment, resulting in a total of 144 pots. After one year, we unpotted the plants and manually separated the roots of each plant individually. Shoots and roots were dried in an oven (72°C for 48 hours) and weighted separately. For each plant, we measured the aboveground, belowground and total dry biomas. Root-shoot dry biomass ratio was then calculated.

Using the setup described above also aimed to answer question (3) does relatedness between individuals influence the diversity of the associated CMN? To do so, the roots from plants grown in well-watered pots were used for root associated fungal community profiling. Because water deficit may influence the development of CMN, the fungal community was only characterized in the well-watered treatment. Total genomic DNA was extracted from approx. 100 mg of root samples using the DNeasy Plant Mini Kit (Qiagen, Germany), according to the kit manufacturer’s instructions. DNA samples were diluted to 3.5 ng/µl after concentration measurement by fluorescence (Quant-IT^TM^Picogreen, Invitrogen, Oregon, USA) and purified by Ampure XP beads, (Agencourt, Beckman Coulter, USA) to eliminate PCR inhibitors. PCR amplification was performed using different primer tag combinations and specific primer sets targeting the ITS2 region (ITS86F: GTG AAT CAT CGA ATC TTT GAA; ITS4: TCC TCC GCT TAT TGA TAT GC; White et al. 1990; Turenne et al. 1999) in 25 μl of reaction mixture as described in Petrolli et al. (2021). Additional negative controls were amplified at this step. After pooling of triplicate PCR reactions, the quality of amplification pools was checked on agarose gels (5 µL, 2 %, 100 V, 20 minutes). Fungal PCR products were purified twice by Ampure XP beads (Agencourt, Beckman Coulter, USA) and quantified by Qubit ® dsDNA HS Assay Kit (Invitrogen, USA). After checking DNA concentration (Quant-IT^TM^ Picogreen, Invitrogen, Oregon, USA), PCR products were pooled together in an equimolar amount of 25 ng. The sample pool was purified twice by Ampure XP beads (Agencourt, Beckman Coulter, USA), MetaFast library preparation and sequencing were performed on an Illumina 2 × 250 MiSeq platform by Fasteris SA (Switzerland).

A pipeline based on VSEARCH (Rognes et al. 2016) and available in GitHub (https://github.com/BPerezLamarque/Scripts/) was used for data processing and the retained fungal annotated reads were subsequently clustered into operational taxonomic units (OTUs) at 97% of identity (Petrolli et al. 2021; Perez-Lamarque et al. 2022). The taxonomic assignment of the OTUs was performed by using UNITE (v.8.0). The OTUs were filtered based on comparison with PCR negative control using the R package DECONTAM, (*prevalence* method) (Davis et al. 2018), fungal kingdom affiliation and their abundance (≥ 10 reads). We used FUNGUILD (Zanne et al., 2019) to classify each OTU according to its functional guilds, and to select OTUs assigned to ectomycorrhizal and endomycorrhizal fungi, after validation by author’s expertise. Then both OTUs tables were normalized to the total number of reads per sample. Bray-Curtis dissimilarity between samples was calculated based on normalized OTU tables for beta-diversity analysis at the OTU level. We performed permutational multivariate analysis of variance (PERMANOVA) in R with the vegan package (Oksanen et al., 2019), by using the capscale() function, to quantify the explained variance by the factors (i.e. kin treatment and families: A1, A2, A3 and A4) (Supplementary Table S1). A p-value of less than 0.05 was considered as significant (with 999 permutations). The shared fungal community was defined by cooccurrence of OTUs associated with two individuals grown in the same pot.

Experiment #2 Nutrient uptake

This second experiment aimed to answer the question: (4) is root nutrient uptake affected by neighbour’s identity? To do so, we use the common experimental setup with the kinship treatment (explained above in the Experimental setup section). In this experiment, each combination of kinship treatment * four family was replicated nine times, resulting in a total of 72 pots. After one year of growth we estimated for each plant nutrient uptake thanks to root nitrate influx. At a physiological level, root nitrogen influx relies on activities of root transport systems and here correspond to the measurement of active transport for nitrate uptake for a given amount of time (here 15 min). Influx was determined using a labelling solution with 15N at three external nitrate concentrations under greenhouse conditions: 30, 300 and 1000 µM. Three replicate pots were used for each concentration. Plants were harvested from pot and roots were gently rinsed with tap water before bathing for 5min in CaCl_2_ to remove nitrogen from root surface. Before ongoing labelling treatment, roots were immersed in a solution with the KNO_3_ concentration for 15 min, to equilibrate nitrate concentration between nutrient solution and apoplast. After acclimation roots were transferred into labelling K15NO_3_ solution for 15 min. After labelling, roots were rinsed into two successive baths with unlabelled KNO_3_ for 5 seconds and 2 minutes respectively to desorb 15N from the root surface. Durations of acclimation, labelling and desorption were based on previous works by Kronzucker et al., (1995) and Fernandez (2019). Shoots, main and fine roots of each individual were then harvested and dried separately at 72°C for 48 hours. We measured the same biomass variables as in experiment#1 and the dry mass of root hair (carefully harvested from each individual) needed for the influx calculation. After determining the respective dry weight, they were pooled and ground to a fine powder (with a ball mill) and micro-weighted. Finally, nitrogen content %N_total_(% of dry weight) and 15N atomic abundance (Q^15^N)were measured in each sample using an Isotope Ratio Mass Spectrometer (IRMS) at SilvaTech facility, INRAE Champenoux. The total amount of N taken up (g) per plant is calculated as follows:

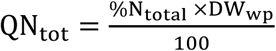

where, %N_total_ is the N content (expressed as % of DW) and DW_wp_ is dry weight of the whole plant (g).

The amount of 15N (g) in each plant was calculated as follow:

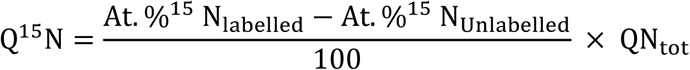

where At. %^15^ N_labelled_ is isotopic abundance of the sample calculated as follow:

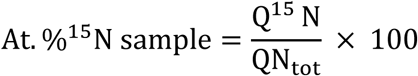

And At. %^15^ N_unlabelled_ is the natural abundance of 15N found in unlabelled sample (mean measured in our unlabelled samples from experiment #1 = 0.3663).

Influx (I) per individual (µmol. g-1. h-1) was calculated as follows:

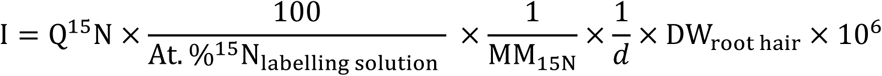

with At. %^15^N_labelling solution_, atomic abundance of 15N (%N) in labelling solution; MM15N, molar mass of 15N (15 g. mole-1), d, duration of the influx measurement (15 minutes) and DW_root hair_the dry mass of root hair (g).

### Statistical analyses

All statistical analyses were performed using R (R ver. 4.0.3). Biomasses variables and output from the IRMS variables were tested for normality and homoscedasticity, using the Shapiro-Wilk and Levene tests. To achieve residual normality, response variables were either log or square root transformed before fitting the model. Variables were back transformed for figures. We compared the treatment effects using two-way ANOVA. Interactions were assessed using type-III tests and multi-comparison test using the Bonferroni post hoc test. When samples size was too low and did not verify normality, we used nonparametric Kruskall-Wallis tests. Unless specified, in all statistical analyses, p < 0.05 was considered statistically significant (α-risk). All data are expressed as mean ±SE.

In experiment #1, we used linear models to test the effect of family, kinship treatment (Kin, Stranger, Control), family (A1, A2, A3, A4), and water deficit treatment (water deficit or well-watered) on aboveground, belowground, and total biomass and on root-shoot ratio. To test whether our plants were in a competitive situation in the pots, we compared the total biomass of the single plant in the Control treatment versus the pooled biomass of the four plants in the Kin and the Stranger treatments using a Kruskall-Wallis test. To ensure that all families have similar inherent growth capacity, we compared individuals in the Control treatment of each family with a Kruskall-Wallis test. In order to assess the competitive ability of each family (i.e. how individuals from a given family perform when growing with individuals from other families), we compared for each of the four families the mean total dry biomasses of individuals grown in the Stranger treatment only. As the position of the plant in the pot (i.e. focal versus surrounding) did not show any significant effect, we tested the Kinship treatment by integrating all plants of the Kin and the Stranger treatments in our model. In experiment #2, we used linear models to test the effect of family and kinship treatment (Kin, Stranger) on biomass variables. We analysed the influx using two-way Anova for each concentration separately. For both experiment#1 and #2, interactions between family and kinship or stress treatments was significant and thus, we analysed the treatment (kinship or stress) for each family separately.

## Results

### Experiment#1

#### Plant response to water-deficit, and kinship and interactions effects

The effect of the kinship treatment, family, water condition, and their interactions on shoot, root, total biomass and the root-shoot ratio are listed in Table 1. Water deficit had a significant negative impact on total dry biomass per pot in the three treatments (Kin, Stranger and Control; Fig.1). Moreover, plants biomass in the Kin and the Stranger treatments was significantly lower total biomass than that from Control treatment in both the water deficit and well-watered condition (p.value < 0.001; Fig.1). Overall, there was no difference in total dry biomass between individuals in the Kin and the Stranger treatment in either water deficit (p.value = 0.9951) and well-watered condition (p.value = 0.9139). Additionally, the total dry biomass of the single plant in the Control treatment (in each water treatment) was not significantly different from the total dry biomass per pot in the Kin and the Stranger treatments (mean Control = 10.98±0.47; mean Kin = 10.65±0.35; mean Stranger = 10.52±0.28; KW p.value = 0.96; data not shown). Moreover, there was no significant differences in the total dry biomass of individual plants grown in the Control treatment between families (p.value = 0.0857) Overall, this result shows that the different families have similar intrinsic growth capacities.

**Figure 1.**
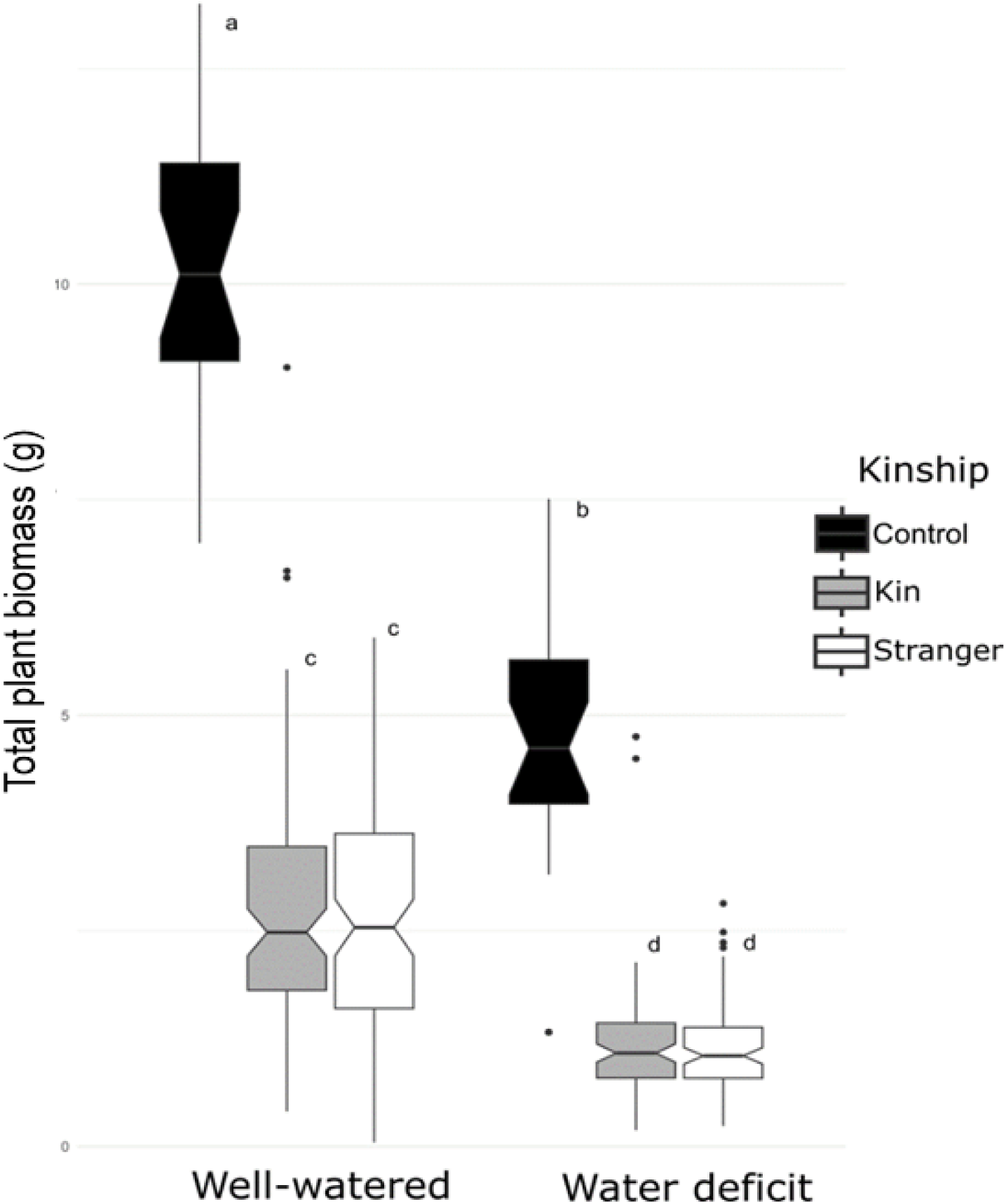
Experiment# 1. Total biomass of plants grown either alone in pot (Control), or with related plants (Kin) or unrelated plants (Stranger), in the water deficit and well-watered conditions. Treatments sharing the same letter are not significantly different for α = 0.05.

**Table 1.** Experiment#1. Results of linear models analysis conducted on all individuals for the different variables in response to the kinship treatment (K), family (F), water condition (W) and their interactions. Significant effects are represented in bold characters.

| *Treatment* | *Total biomass* |  |  | *Shoot biomass* |  | *Root biomass* |  | *Root-shoot ratio* |  |
| --- | --- | --- | --- | --- | --- | --- | --- | --- | --- |
|  | *Df* | *F-stat* | *p.value* | *F-stat* | *p.value* | *F-stat* | *p.value* | *F-stat* | *p.value* |
| *K* | 1 | 0.603 | 0.438 | 0.584 | 0.445 | 0.605 | 0.437362 | 0.008 | 0.93020 |
| *F* | 3 | 8.757 | <**0,001** | 10.546 | <**0,001** | 6.978 | <**0,001** | 4.475 | <**0,001** |
| *W* | 1 | 218.240 | <**0,001** | 227.814 | <**0,001** | 179.145 | <**0,001** | 2.431 | 0.11984 |
| *K\*F* | 3 | 9.291 | <**0,001** | 11.134 | <**0,001** | 6.694 | <**0,001** | 1.365 | 0.25334 |
| *K\*W* | 1 | 0.026 | 0.872 | 1.070 | 0.302 | 0.215 | 0.643336 | 5.817 | **0.01638** |
| *F\*W* | 3 | 1.367 | 0.253 | 1.762 | 0.154 | 1.321 | 0.267380 | 2.716 | **0.04465** |
| *K\*F\*W* | 3 | 0.660 | 0.577 | 0.383 | 0.765 | 0.693 | 0.556714 | 0.251 | 0.86073 |
| *Residuals* | 355 |  |  |  |  |  |  |  |  |

Water-deficit had a global negative impact on total, shoot and root biomasses but not on the root-shoot ratio. Kinship had no significant impact on biomasses and root-shoot ratio. The interaction between the water condition was significant with the kinship and family only for the root-shoot ratio (p.value = 0.016 and 0.045 respectively). Family was significant for all variables. In addition, the three-way interaction was never significant. We found that the family effect interacts significantly with the kinship for all biomass variables except root-shoot ratio, and with the water condition for the root-shoot ratio only. Thus, we analysed the data for each family separately.

#### Family competitive ability

For each family, we tested for competitive ability by summing up and comparing the total dry biomass of plants from different pots where they were grown only with strangers (Stranger treatment). When grown with strangers, we found that individual from the A1 family exhibit less total dry biomass than the A2 and A4 families (p.value = 0.0025 and <0.001 respectively). In contrary, individuals from the A2 family performed better than A1 and A3 (p.value <0.001 in both case) but showed no difference with the A4 family (p.value = 0.93). A similar result (data not shown) was found for plants under water-deficit condition.

#### Kinship effect for each family separately

For each family, water condition did not interact significantly with the kinship treatment for any variable (not shown). Thus, for the following results, we only considered individuals grown in the well-watered condition and without plants from the Control treatment. Plants from the A1 and A2 family showed a kinship effect (Tab. 2). However, the A1 and A2 families showed opposite results. Individuals from the A1 family grown in the Kin treatment had significantly higher total biomass (mean_kin_ = 3.00±0.26; mean_stranger_ = 1.77±0.18), shoot biomass (mean_kin_ = 1.29±0.11; mean_stranger_ = 0.68±0.07), and root biomass (mean_kin_ = 1.71±0.18; mean_stranger_ = 1.08±0.12) than individuals grown in the Stranger treatment (Fig.2 and Table 2). Conversely, individuals from the A2 family grown in the Stranger treatment had significantly higher total biomass (mean_kin_ = 2.48±0.27; mean_stranger_ = 3.08±0.28), shoot biomass (mean_kin_ = 1.08±0.11; mean_stranger_ = 1.25±0.12), and root biomass (mean_kin_ = 1.40±0.16; mean_stranger_ = 1.82±0.18) than individuals grown in the Kin treatment (Fig.2 and Tab.2). Individuals from the A3 and A4 families did not show any effect of the kinship treatment (Tab. 2).

**Figure 2.**
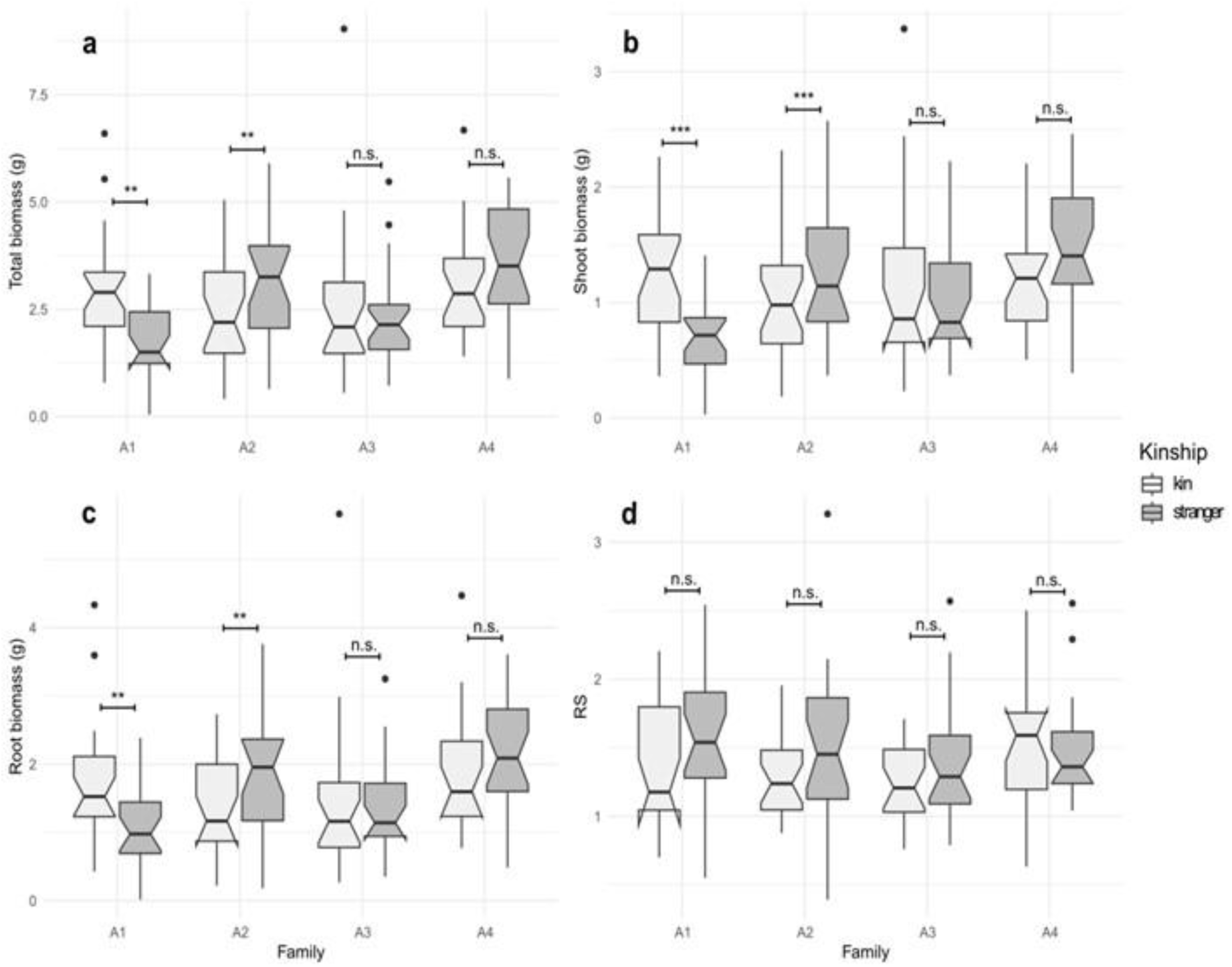
Experiment# 1. Boxplot for (a) total biomass, (b) shoot biomass, (c) root biomass (all in grams) and (d) root-shoot ratio (RS), for each family in the Kin and Stranger treatments from well-watered condition. **, significant difference for α <0.01; ***, significant difference for α <0.001 ; n.s., non-significant.

**Figure 3.**
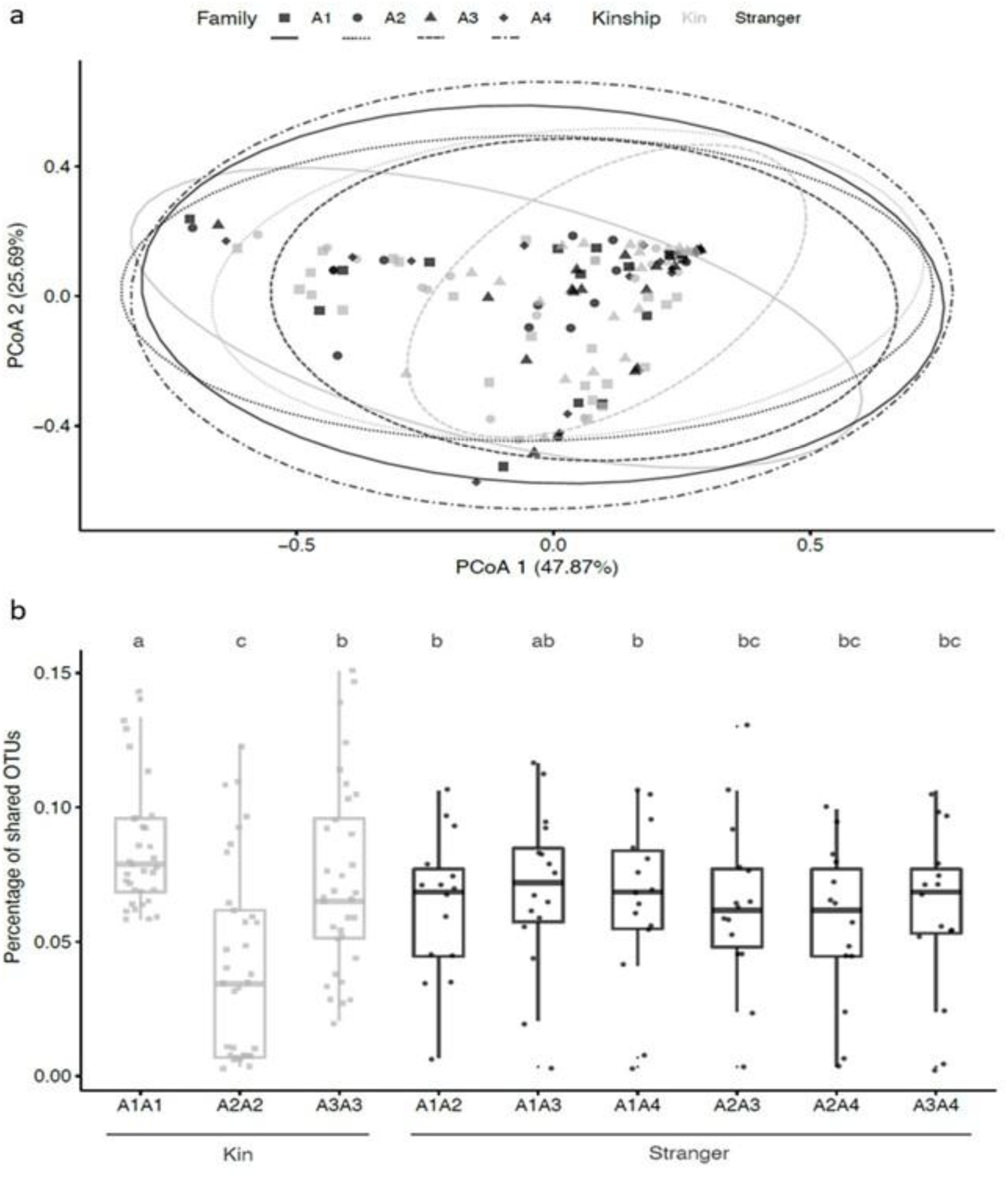
Exp#l. Effect of the kinship treatment and family on the total and shared ectomycorrhizal fungal community. (a) PCoA based on Bray-Curtis dissimilarity between Allier root samples belonging to four families (Al. A2. A3. A4) in the Kin and Stranger treatment, (b) Percentage of ectomycorrhizal shared OTUs between individuals in the Kin and Stranger treatments. The percentage of shared OTUs between two plants is relative to the total number of ectomycorrhizal fungi. Lowercase letters indicate significant differences for Kruskall-Wallis test (p.value <0.05). We have specified the family combinations in part B of the figure.

**Table 2.** Experiment#1. Effect of the kinship treatment on total, shoot and root biomass and root-shoot ratio for each family separately. n.s. non-significant; K, kin treatment; S, stranger treatment. Significant p.values are represented in bold characters below the differences between the Kin and Stranger treatments.

| *Family* | *Total biomass* | *Shoot* | *Root* | *RS* |
| --- | --- | --- | --- | --- |
| *All* | n.s. | n.s. | n.s. | K > S<br>**0.029** |
| *A1* | K > S<br>**0.002** | K > S<br>**<0.001** | K > S<br>**0.004** | n.s. |
| *A2* | S > K<br>**0.002** | S > K<br>**<0.001** | S > K<br>**0.009** | n.s. |
| *A3* | n.s. | n.s. | n.s. | n.s. |
| *A4* | n.s. | n.s. | n.s. | n.s. |

#### Root associated fungal community according to kinship and family

Overall, family has a significant impact on the ectomycorrhizal and endomycorrhizal fungal communities in root samples (p.value = 0.028 in both case; Tab. 3), while the kinship treatment had a significant impact on endomycorrhizal community only (p.value = 0.006; Tab. 3). For the ectomycorrhizal fungi, samples belonging to the A3 family clustered (on the 1^st^ axis), as compared to the other samples (Fig. S1A). Two different trends were observed for the A1 and A2 families, regarding the ectomycorrhizal shared community. For A1, the percentage of ectomycorrhizal shared OTUs between two individuals grown in the kin treatment (A1A1 on Fig.S1B) was significantly higher (p.value <0.05) than in the Stranger treatment compared to A1A2 and A1A4 but not when compared to A1A3 (Fig. S1B). Conversely, A2 family showed opposite results with a greater percentage of ectomycorrhizal shared OTUs in the Stranger treatment compared to the Kin treatment. Similarly, the percentage of the endomycorrhizal shared community was greater in the Stranger treatment than in the Kin treatment, for the A2 family (Fig. S1B).

**Table 3.** Experiment#1. Percent of variance explained by the family (F), the kin treatment (K) and their interaction on diversity (Beta-diversity Bray-curtis matrice) of endo- and ectomycorrhizal fungi (permanova), model ∼ Family * Kinship. Significant effects are represented in bold characters.

|  | *Ectomycorrhizal fungi* |  |  | *Endomycorrhizal fungi* |  |  |
| --- | --- | --- | --- | --- | --- | --- |
|  | *Df* | *F-stat* | *p.value* | *Df* | *F-stat* | *p.value* |
| *F* | 3 | 1.8955 | **0.028** | 3 | 1.4012 | **0.028** |
| *K* | 1 | 0.7156 | 0.553 | 2 | 1.8873 | **0.006** |
| *K\*F* | 2 | 1.2081 | 0.265 | 2 | 1.5694 | **0.018** |
| *Residuals* | 121 |  |  | 83 |  |  |

### Experiment#2

#### Influx

Similarly, to experiment#1, we control for the competitive ability by comparing the mean influx (i.e. nitrogen uptake) for individuals from a given family in the Stranger treatment. None of the four families showed differences in competitive ability (Fig S2).

Overall, the kinship treatment had a significant impact on influx at 30µmol and 300µmol and family had a significant effect on the influx only at the 300µmol concentration (Supplementary material table S2). The interaction between the kinship treatment and family was not significant at any concentration (Table S2). However, considering our results from experiment #1 that showed a strong interaction between the kinship treatment and family we then analysed the influx at for each family for the three concentration separately. This allows us to show that at 30µmol, individuals from the A3 family exhibits a significantly higher influx when grown in the Kin condition (mean_kin_ = 3.162±0.45; mean_stranger_ = 1.783±0.42; p.value = 0.012; Fig.4). The same tendency was observed for the A2 family (mean_kin_ = 2.102± 0.226; mean_stranger_ = 1.682±0.506; Fig. 4) although this result was only marginally significant (p.value = 0.057).

**Figure 4.**
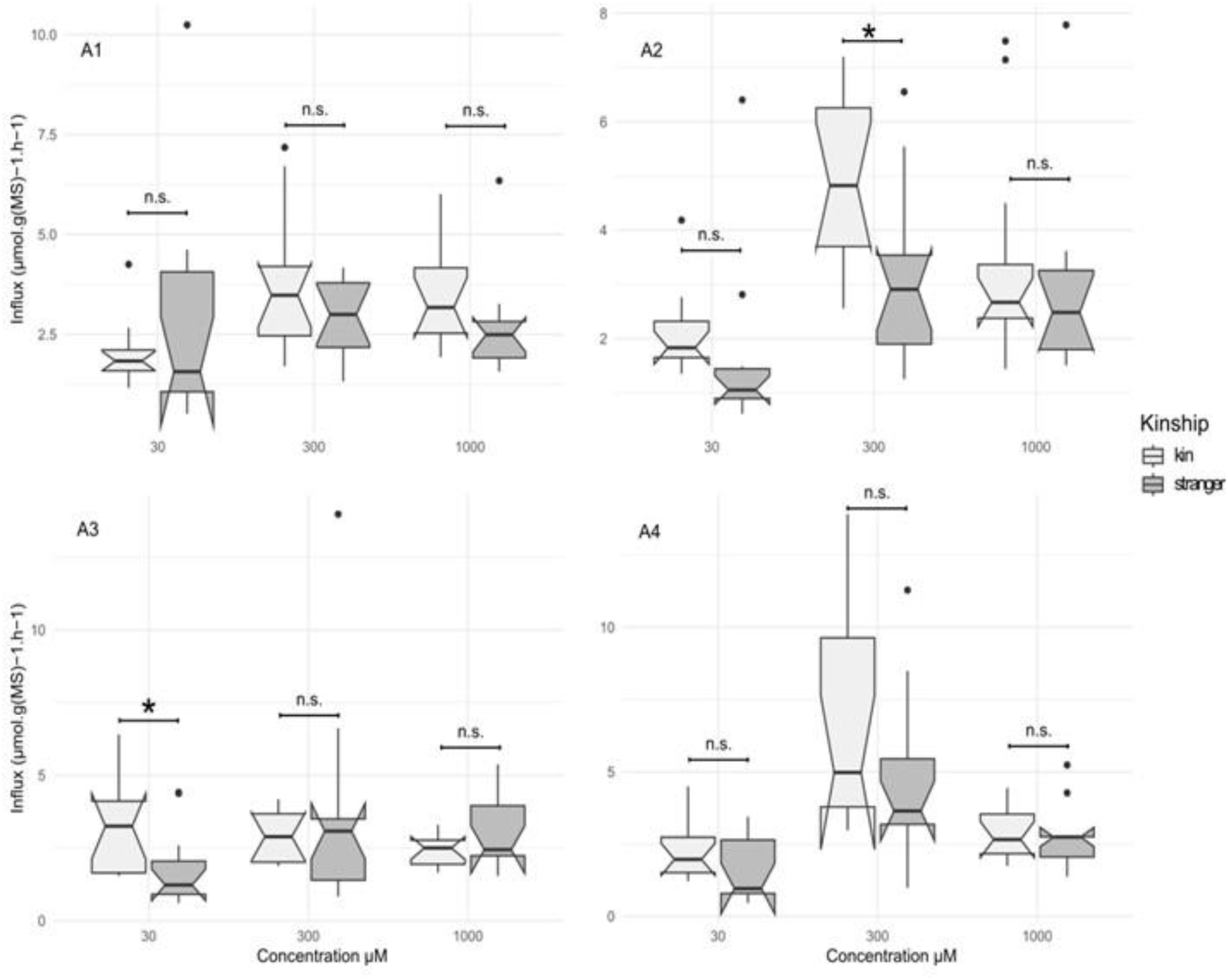
Exp#2. Influx in the two kinship treatments and the three concentrations for each family separately (Al, A2, A3 and A4). *, significant difference with a p.value <0.05; n.s., non-significant

At 300µmol, of the four families, only individuals from the A2 family grown in the Kin treatment showed a significantly higher influx than those grown in the Stranger treatment (mean_kin_ = 4.806± 0.453; mean_stranger_ = 3.083± 0.488, anova p.value = 0.010; Fig.4).

## Discussion

### Competitive ability overrides kinship effect in plant growth and nutrient uptake

The phenotypic response to kinship treatment differed among families both in amplitude and direction. In particular, the A1 and A2 families showed opposite biomass trends. Individuals from the A1 family grown in the Kin treatment exhibit higher biomass (total, shoot and root) than those grown in the Stranger treatment, whereas the A2 family displayed the inverse trend. At first sight, these opposite results could be interpreted with the two commonly frameworks used in kin studies in plants. The kin selection framework states that individuals growing with relatives should invest less in competitive traits to increasing their inclusive fitness (Ehlers and Bilde 2019) and this accounts for the features of family A2. In plants, reduced root biomass is considered a cooperative trait (i.e. a positive interaction) as it leaves more space and thus more resources available for neighbours (File et al. 2012a; Bilas et al. 2021). By contrast, when plants grown with related individuals exhibit greater belowground biomass than plants grown within non-related individuals, this is mostly interpreted with the niche partitioning framework (File et al. 2012a; Subrahmaniam et al. 2021) and this may account for the features of family A1. Niche partitioning predicts that related individuals exhibit small niche differentiation and thus overlap more in their niche use, leading to a greater competition with each other compared with unrelated conspecifics (Young, 1981, Milla et al. 2009, Ehlers and Bilde, 2019). Moreover, investment in root biomass in soil with limited resources is commonly understood as a competitive behaviour (i.e. negative interaction) because an individual will have greater belowground colonization and nutrient uptake (Anten and Chen, 2021).

Kin selection and niche partitioning frameworks could occur simultaneously, possibly acting on different traits and thus different niche components (File et al. 2012a) and may cause complications in the interpretation of experimental results (Anten and Chen, 2021). Another confounding factor is intrinsic competitive ability, i.e. the relative growth of individuals grown in competition compared to grown alone (Masclaux et al. 2010; Zhang et al. 2019). Here, individuals from the A1 family invested more growth when growing with kin, thus showing a competitive behaviour. However, individuals from the A1 family had lower biomasses than individuals from most other families (A2, A4) in the Stranger treatment, suggesting that they are poor competitors. Thus, when individuals from the A1 family were grown together in the Kin treatment, they performed better than in the Stranger treatment, where they experience stronger competition. Conversely, individuals from the A2 family are considered good competitors as they grew better than all the other families in the Stranger treatment. When grown with their relatives, they suffer from a stronger competition than when in the Stranger treatment, with less competitive individuals. As such, the most appropriate and parsimonious interpretation for our results is therefore differential competitive abilities between families instead of kin selection or niche partitioning frameworks. This interpretation is also valid for the two families (A3 and A4) that showed no differences in biomass in relation to the kinship treatment; competitive ability could then be a polymorphic trait (i.e. trade-off between competition depending on the identity of the neighbours) and vary at an intraspecific level. Yet, only few studies have considered the competitive ability of the families used in their kin studies designed as a relevant interpretation. To our best knowledge, it has only been taken into consideration in two studies *Arabidopsis thaliana* (Masclaux et al. 2010) and *Medicago minima* (Tomiolo et al. 2021), but remains overlooked in other kin studies (Mazal et al. 2023).

Variances of biomass among plants grown in the same pots in the Kin and the Stranger treatment could also help identifying differences in competitive ability (Tomiolo et al. 2021). One hypothesis is that in the Kin treatment, biomasses will have a lower variance suggesting a weaker competition among related individuals. Conversely, in the Stranger treatment, higher variances for biomasses are expected, indicating a stronger competition (Ehlers and Bilde 2019). In this last case, when growing in pots, individuals that would exhibit higher biomass compared to the other plants in the pot, would be considered to express a better competitive ability. In our case, within pot variances did not differ between the Kin and the Stranger treatment (data not shown). Following Fréville et al. 2019, we emphasize the need to consider competitive ability in future kin recognition and selection studies, to avoid misleading conclusions on kin recognition.

We detected no kinship effect when considering nutrient uptake at higher ammonium concentration (1000 µmol). This could indicate that at 1000 µmol, ammonium was abundant and thus not limiting, allowing all individuals to draw on nutrients and hiding potential kinship effects. Interestingly, for the two lowest concentration (30 and 300 µmol), individuals grown in the Kin treatment showed a higher nitrogen uptake than those grown in the Stranger treatment. However, since we did not find differences in competitive ability between families, we cannot explain this result by a difference in competitive ability as for biomasses. Our results suggest that individuals could be able to detect related individuals and increase their nutrient uptake. However, they are difficult to interpret because our data on biomasses does not indicate kin discrimination, and thus kin recognition. Nevertheless, if individuals are capable of kin recognition through nutrient uptake, the increase in their nitrogen absorption could represent an increased competition. It is then difficult to conceive that individual detect other related individuals in order to increase competition with them. The associated cost to kin recognition mechanisms would not be counterbalanced by any increase competition between relatives. Thus, our results do not fit in the kin recognition framework. Alternatively, they could be explained by the niche partition hypothesis, because in this case competition is stronger when individuals grow with related individuals. Nutrient absorption occurs at the fine root parts of the plant *via* specific transporters that are coded by multigenic families (Le Deunff et al. 2016). It is thus expected that closely related individuals share the same multigenic families and thus exhibit similar nutrient uptake capacities. However, these results are open to interpretation because influx was measured over a short period of time (40 minutes) compared to the duration of the experiment (one year). Moreover, we cannot exclude that the modification of mycorrhizal community, as discussed below interfere in the nutrient uptake capacities.

### Drought did not change the outcome of the interaction

Water deficit was significant in our experiment, since individuals experiencing water deficit were significantly smaller than those from well-watered condition. Strikingly, for each of the four family, the water condition did not change the outcomes of interaction between individuals, suggesting a robust pattern. Similarly, Goddard et al. (2021) reported that water limitation had no impact on the results of the kinship interactions in *Glechoma hederacea*. The stress gradient hypothesis (Bertness and Callaway 1994, Callaway et al. 2002) predicts that competition is more common at low and high level of stress. Black poplar individuals exhibit a strong water-deficit tolerant strategy (Corenblit et al. 2014) and thus interactions between individuals that exhibit a similar stress-tolerant strategy could be negative at both end of the stress gradient hypothesis especially when resources are limited (Maestre et al. 2009). Moreover, when a stress factor is added to a resource stress, negative interactions should be prevalent (Adams et al. 2021). Thus, in our case, the water-deficit may have been too intense, and added to the competition ongoing in a limited space (i.e. the pots) leading to excessive stress and thus competition was the predominant interaction we observed.

Other types of environmental constraints (e.g. nutrient deficit; shear stress; sediment burial) could also be tested to search for a change in the type of interactions. For example, several studies have found more conclusive results using a nutrient limitation. In cultivars of *Pisum sativum* the cooperative response toward kin plants increases with nutrient limitation (Pezzola et al. 2020), while interactions among *Setaria itlica* individuals at different nutrient conditions are influenced by both neighbour’s identity and density (Husain Jaafry et al. 2020). In another case, the stress gradient hypothesis could predicts a change in the interaction type in relation to stress (Till-Bottraud et al. 2012). In *Nothofagus pumilio* case, environmental constraints play a crucial role in multi-stemmed tree clusters at edge of secondary growth forests stressed by strong winds. At these edges where winds are intense, *N. pumilio* seedlings survived significantly better in clusters than alone (McIntire and Fajardo 2011). Conversely, no merged trees were observed within the forest where winds are weaker, and where seedlings survived better isolated than in clusters. Interestingly, the different stems of merged trees were often strongly related (half-sibs, full-sibs or even higher relatedness), indicating that kin selection might favour the interaction. Thus, these results lead us to assume that kin recognition and discrimination may only be expressed under certain ecological conditions which emphasizes the value of including different types of stress in kin studies.

### Effect of kinship on root fungal communities

We showed a significant effect of the plant family on both endo- and ectomycorrhizal diversity found in black poplars (Table 3). Intraspecific polymorphism for this symbiosis, although overlooked, is expected from several works (Johnson et al. 2012). In fact, plant intraspecific variability can lead to variable fungal patterns in the root’s proximity (Goldmann et al. 2020) and can have significant positive effects on plant-fungal interactions (Hazard et al. 2017). In *Populus* trees, microbial communities differ across tissue and within leaf, endosphere fungal communities are strongly impacted by the host genotype (Cregger et al. 2018). Although the authors found different results between different *Populus* species (*Populus deltoides* and *P. deltoides x trichocarpa*) our results suggest that intraspecific diversity can also be a driver of mycorrhizal community assemblage. Thus, the establishment of a common mycorrhizal network could be influenced by plant genotypes and identity of neighbours and could contribute to the differences in growth found for these two families.

Moreover, the percentage of shared ectomycorrhizal OTUs between individuals displayed opposite tendencies in relation to the families considered. For the A1 family, we observed that individuals in the Kin treatment shared more OTUs with their neighbours than individuals in the Stranger treatment, whereas the opposite is observed for the A2 family (Fig. S3). We have previously showed that A1 and A2 families have both a trend to exhibit the highest individual total biomass in condition where the percentage of shared ectomycorrhizal OTUs was the highest (respectively the Kin and the Stranger treatment). In this particular case sharing mycorrhizal fungi and their costs could contributes positively to growth, as already described in some CMN interactions (Selosse et al. 2006; van der Heijden et al. 2015; Karst et al. 2023). Although we did not investigate the establishment of CMN, our results can be viewed in the context of those from other studies. In fact, several studies on kin recognition in plants investigated the role of CMN on plant growth and nutrient uptake. File et al. (2012b) show that related *Ambrosia artemisiifolia* individuals, benefited more from the presence of a common mycorrhizal network, in terms of phosphate uptake and suppression of pathogens, than non-related individuals. The authors argued that the CMN can be involved in kin recognition clues, whereby plants recognized the presence of kin through exudates or through secondary metabolites released by neighbouring plants and transported through the CMN, which induce them to invest resources in the network. Investment in a CMN is a cooperative behaviour in which the costs associated with investment in a common network are smaller than the accumulated benefit because it increases the resource availability (Anten & Chen 2021). Pickles et al. (2017), using ^13^C as a tracer to observe exchanges between individuals via mycorrhizal networks, showed that ^13^C concentrations in mycorrhizal biomass as well as in ectomycorrhizal plants were higher in pairs of related individuals than in pairs of unrelated individuals. Furthermore, the transfer of carbon compounds between individuals was significantly higher in related individuals. Thus, plants may be able to selectively transfer nutrients to related individuals (Pickles et al. 2017). The study of the mycorrhizal community in kin recognition studies appears to be a promising avenue for understanding the underlying mechanisms of kin recognition and the outcomes of interactions between related individuals.

## Conclusion

The present study investigated the effect of neighbour relatedness and water deficit on intraspecific interaction. Young *P. nigra* kin groups studied revealed variable plastic responses, namely for the growth variables, to neighbour identity and to a water deficit. A main result of our study was the fact that differences in biomasses were explained by differences in competitive ability and not by the usual frameworks used in kin studies (i.e. kin recognition and niche partitioning) and could potentially be linked with higher mycorrhizal fungal sharing. We showed a significant effect of the plant family on both endo-and ectomycorrhizal diversity, suggesting that the study of CMN could be a relevant factor in the understanding of the mechanism of kin recognition in plants. Moreover, higher nutrient absorption among individuals who grew up in the Kin treatment indicates a stronger competition between relatives for nitrogen uptake and thus suggest the niche partitioning hypothesis for this trait. These results emphasise the fact that depending on the studied traits, different frameworks can be called simultaneously to interpret results. Thus, results obtained when growing plants in competition with related individuals should be carefully analysed. We advocate to investigate and test for confounding factors, especially the competitive ability of kin groups which is rarely assessed, before wrongly conclude to the evidence of kin recognition or discrimination in plants.

## Acknowledgments

The author(s) would like to thank SILVATECH (Silvatech, INRAE, 2018. Structural and functional analysis of tree and wood Facility, doi: 10.15454/1.5572400113627854E12) from UMR 1434 SILVA, 1136 IAM, 1138 BEF and 4370 EA LERMAB from the research center INRAE Grand-Est Nancy for its contribution to isotopic analysis. SilvaTech facility is supported by the French National Research Agency through the Laboratory of Excellence ARBRE (ANR-11-LABX-0002-01).

## Fundings

The first author was funded for his PhD by the French Ministry of National Education, Higher Education and Research. This research was financed by the French government IDEX-ISITE initiative 16-IDEX-0001 (CAP 20-25). M.-A. S and A.B. are supported by grants from the Institut Universitaire de France and anonymous donators.

## Notes

### Competing Interest Statement

The authors have declared no competing interest.

