## Supplementary table and figure for "Kin discrimination in plants: competitive ability over kinship response"

**Supplementary material**

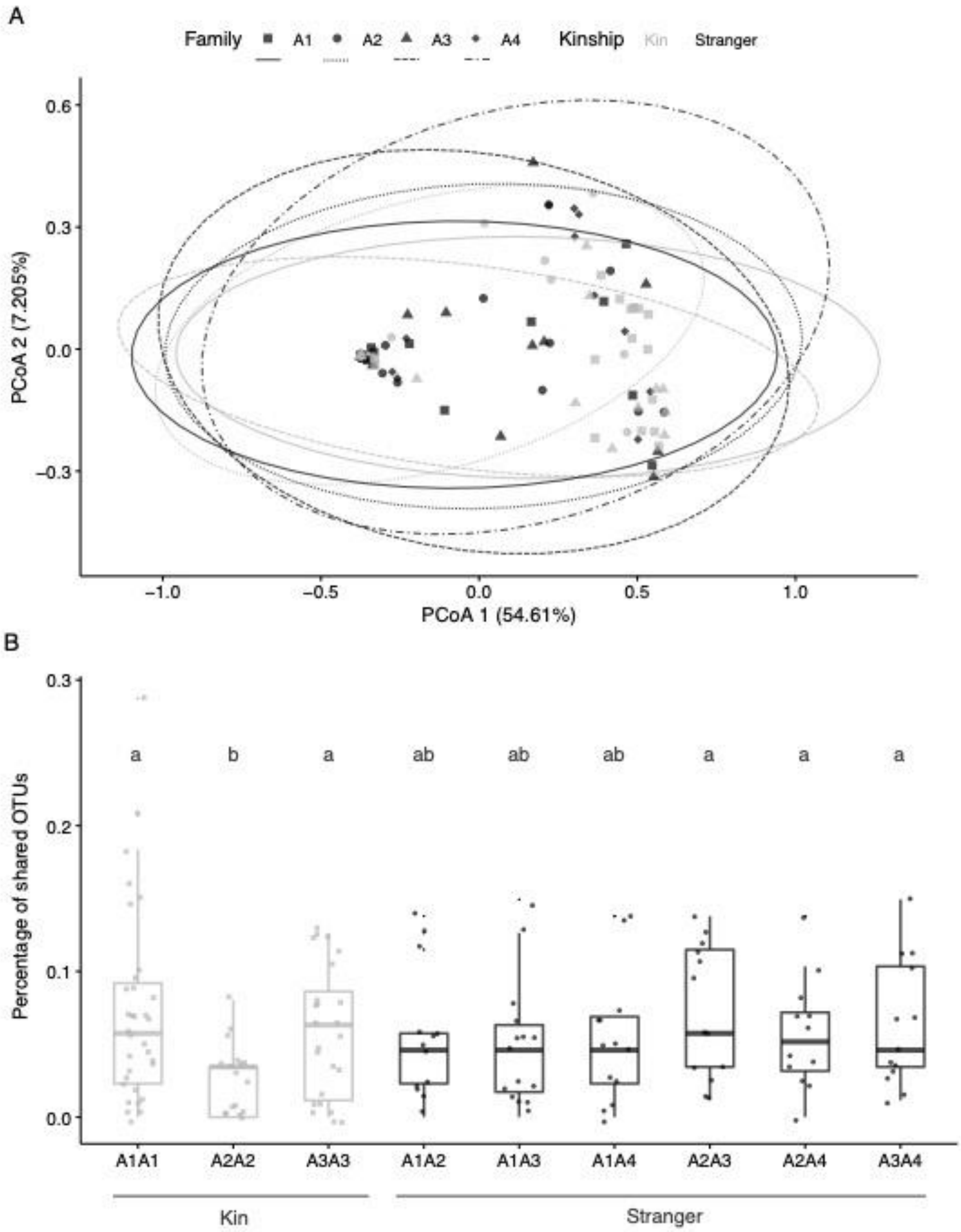


**Figure S1.** Exp#1. Effect of poplar family and kinship on endomycorrhizal fungal community. A) PCoA based on Bray-Curtis dissimilarity between Allier root samples belonging to four families (A1, A2, A3, A4) in the Kin and Stranger treatment. B) Percentage of endomycorrhizal shared OTUs between individuals. The number of shared OTUs between two plants is relative to the total number of endomycorrhizal OTUs. Kruskall-Wallis test was used for statistical analysis (*P* <0.05). Lowercase letters indicate significant differences between kinship pairs.

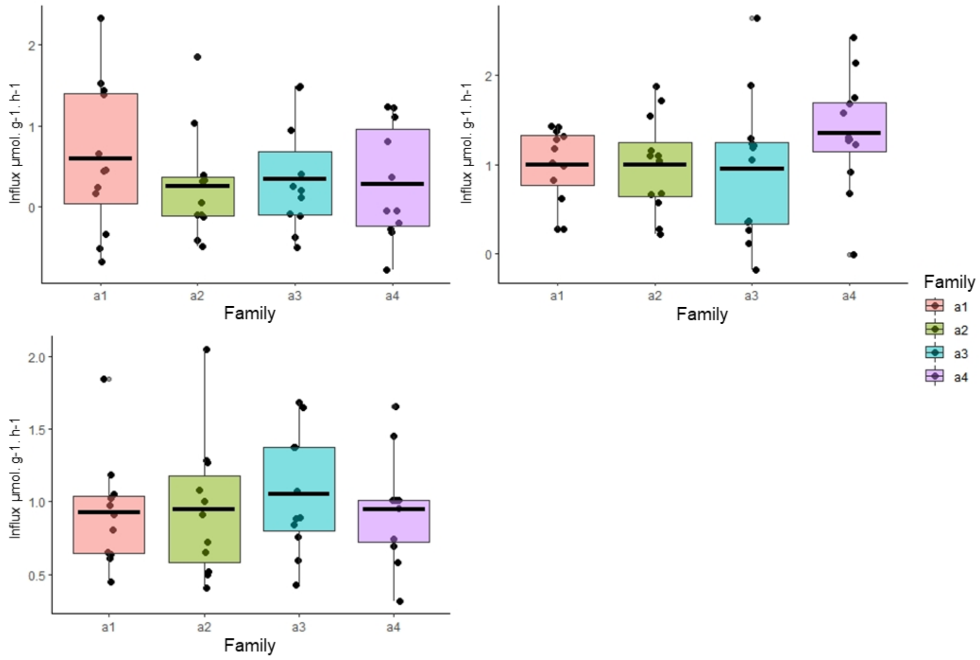


**Figure S2**. Exp#2. Mean influx per family used (A1, A2, A3, A4) measured from individuals grown in the Stranger treatment for each concentration. There were no significant differences between families.

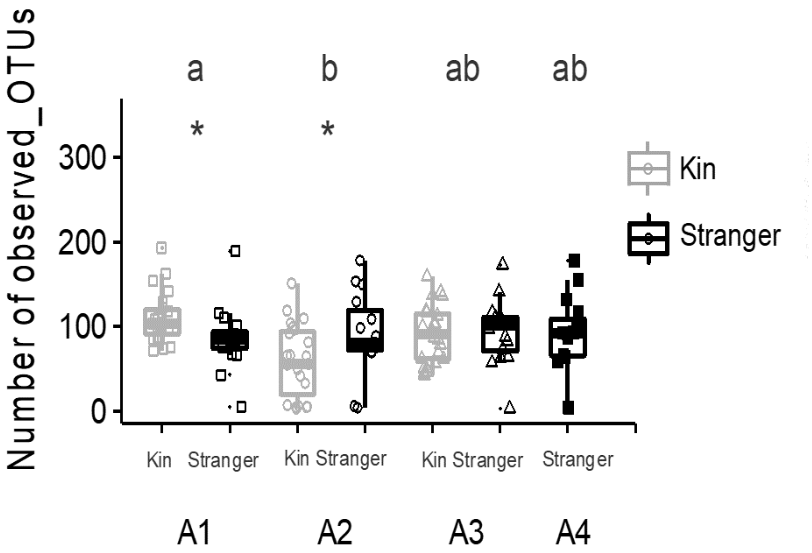


**Figure S3.** Exp#1. Effect of poplar family and kinship on fungal richness (number of fungal OTUs per plant). The Kruskall-Wallis test was used for statistical analysis (*P* <0.05). Lowercase letters and stars indicate significant differences between families and kinship pairs, respectively.

**Table S1.** Explained variance of parameters affecting fungal diversity in Allier population, based on beta-diversity. This includes all plant families and both kinship (PERMANOVA), model ~ Family * Kinship was applied. Significant effects are represented in bold characters.

| | Ectomycorrhizal fungi | | | | Endomycorrhizal fungi | | | |
|---|---|---|---|---|---|---|---|---|
| | *Df* | *F-stat* | *R2 (%)* | *p.value* | *Df* | *F-stat* | *R2 (%)* | *p.value* |
| *Kinship* | 1 | 0.7156 | 0.65 | 0.553 | 2 | 1.8873 | 4.37 | **0.006** |
| *Family* | 3 | 1.8955 | 5.15 | **0.028** | 3 | 1.4012 | 4.87 | **0.028** |
| *Kinship * Family* | 2 | 1.2081 | 2.19 | 0.265 | 2 | 1.5694 | 3.63 | **0.018** |
| *Residuals* | 121 | | 109.6 | | 83 | | 96.06 | |

**Table S2**. Experiment #2. Results of linear models analysis conducted on all individuals for the influx in response to the kinship treatment, family and its interactions. Significant effects are represented in bold characters.

| | | *Influx* | | | | | |
|---|---|---|---|---|---|---|---|
| | | *30µmol* | | *300µmol* | | *1000µmol* | |
| | *Df* | *F* | *p.value* | *F* | *p.value* | *F* | *p.value* |
| *Kinship* | 1 | 10,7592 | **0,001505** | 7,3396 | **0,008139** | 0,6307 | 0,4293 |
| *Family* | 3 | 0,6639 | 0,576552 | 4,7936 | **0,003894** | 0,2801 | 0,8396 |
| *Kinship * Family* | 3 | 1,1194 | 0,34588 | 0,8014 | 0,496438 | 1,348 | 0,2645 |
| *Residuals* | 85 | | | | | | |
